# Sialic Acid Identity Modulates Host Tropism of Sialoglycan-binding Viridans Group Streptococci

**DOI:** 10.1101/2025.06.24.660003

**Authors:** KeAndreya M. Morrison, Rupesh Agarwal, Haley E. Stubbs, Hai Yu, Stefan Ruhl, Xi Chen, Paul M Sullam, Barbara A Bensing, Jeremy C. Smith, T. M. Iverson

**Affiliations:** Department of Pharmacology, School of Graduate Studies, Meharry Medical College, Nashville, TN, USA; UT/ORNL Center for Molecular Biophysics, Oak Ridge National Laboratory, TN, USA; Chemical and Physical Biology Program, Vanderbilt University, Nashville, TN, USA; Department of Chemistry, University of California, Davis, CA, USA; Department of Oral Biology, University at Buffalo, Buffalo, NY, USA; Division of Infectious Diseases, Veterans Affairs Medical Center, Department of Medicine, University of California, San Francisco, and the Northern California Institute for Research and Education, San Francisco, CA, USA; Department of Biochemistry and Cellular and Molecular Biology, University of Tennessee, Knoxville, TN; Departments of Pharmacology and Biochemistry, Vanderbilt University, Nashville, TN, USA

**Author notes:** Current address (Rupesh Agarwal).

**Keywords:** cross-reactivity, host preference, pathogenesis, commensalism, oral microbiome, infective endocarditis, sialic acid, host receptor, adhesin, Siglec-like binding region, streptococci

## Abstract

Microbial interactions with multiple species may expand the range of potential hosts, supporting both pathogen reservoirs and zoonotic spillover. Viridans group streptococci interact with host cells by engaging protein-attached glycosylations capped with terminal sialic acids (sialoglycans). One potential origin for host tropism of these streptococci arises because humans exclusively synthesize the N-acetylneuraminic acid (Neu5Ac) form of sialic acid, while non-human mammals synthesize both Neu5Ac and a hydroxylated N-glycolylneuraminic acid (Neu5Gc). However, the link between binding preference for these sialic acids and preference for host has not been tested experimentally. Here, we investigate sialoglycan-binding by Neu5Ac/Neu5Gc cross-reactive Siglec-like binding regions (SLBRs) from two strains of streptococci, *Streptococcus gordonii* strain Challis (SLBR_Hsa_) and *Streptococcus sanguinis* strain SK36 (SLBR_SrpA_). Structural and computational analyses of SLBR_Hsa_ identified molecular details for the binding of disaccharides capped in Neu5Ac or Neu5Gc. Engineering SLBR_Hsa_ and SLBR_SrpA_ for narrow selectivity to synthetic Neu5Gc-terminated glycans shifted the binding preference from authentic human plasma receptors to plasma receptors from rat sources. However, host receptor preference did not fully recapitulate purified Neu5Ac/Neu5Gc-capped sialoglycan preference. These findings suggest that sialic acid identity modulates, but does not uniquely determine, host preference by these streptococci. This work refines our understanding of host specificity and challenges prevailing assumptions about the relative role of sialic acids in host tropism.

## Introduction

Viridans group streptococci are among the bacteria that can engage sialic acid capped glycans (sialoglycans) on host cells^1^. This host adherence promotes oral colonization when the sialic acid is attached to the glycans on salivary proteins, such as MUC7^2–4^. Host adherence also supports endovascular pathogenesis^5,6^. Indeed, engagement of sialoglycans attached to platelet glycoprotein Ibα^7^ (GPIbα) is among the first committed steps in bacterial infective endocarditis, a serious infection of the heart valves that may result in heart failure, stroke, or permanent valve damage, even with aggressive treatment. Viridans group streptococci are responsible for 40%-60% of bacterial infective endocarditis cases^8,9^. In-hospital mortality for these infections is approximately 10%, while one- and five-year mortality rates are estimated as 22-37% and 37-53%, respectively^10–13^.

Streptococci bind to sialoglycans using an adhesin that contains a domain related to mammalian <u>S</u>ialic acid-binding immuno<u>g</u>lobulin(Ig)-like <u>lec</u>tins (Siglecs)^14^. In streptococci, this binding domain is called the Siglec-like binding region (SLBR)^14^. Both SLBRs and Siglecs are organized around a V-set Ig fold that is predominantly comprised of β-strands and that has a standard nomenclature. The strands of the Ig fold are designated A-G and intervening loops are named based upon the adjacent strands^14^; for example, the AB loop connects the A strand and the B strand. The sialoglycan itself binds above a conserved ΦTRX sequence motif on the F strand, where Φ is a hydrophobic amino acid, most commonly tyrosine.

SLBRs engage sialic acid capped terminal tri- and tetra-saccharides^15,16^ on heterogeneous glycosylations, which can subtly differ in identity and presentation between individuals^17,18^. Importantly, the preferred sialoglycan ligand of SLBRs correlates with disease severity in an animal model, with selective binding to a sialyl-T antigen-capped receptor on GPIbα most strongly promoting endocardial infection^6^. As revealed by chimeragenesis, control over the preferred sialoglycan ligand disproportionately involves direct interactions with side chains on three loops that surround the binding site: that CD loop, the EF loop, and the FG loop^16^. Perhaps surprisingly, it is difficult to predict the sialoglycan binding spectrum of SLBRs. Even SLBRs with >90% identity can exhibit a different sialoglycan binding repertoire^16,19^.

Humans and non-human animals can differ in their production and presentation of sialoglycans. Within the context of complex glycosylations, these differences include the underlying glycan composition, the preferred glycosyl linkages, the local distribution of different sialoglycans in biological niches, and even the sialic acid itself^20,21^. Sialic acids are nine-carbon acidic sugars with more than 50 biological forms^17,21,22^. Whereas humans only synthesize N-acetylneuraminic acid (Neu5Ac, **Fig. 1A**) and its derivatives, most non-human animals can convert Neu5Ac to N-glycolylneuraminic acid (Neu5Gc, **Fig. 1B**)^23^ and its derivatives. Neu5Ac and Neu5Gc only differ by a hydroxyl group appended at the C11 position^24,25^ (**Fig. 1B**). This is subtle, yet some infectious agents can distinguish between the two^26–29^.

**Figure 1.**
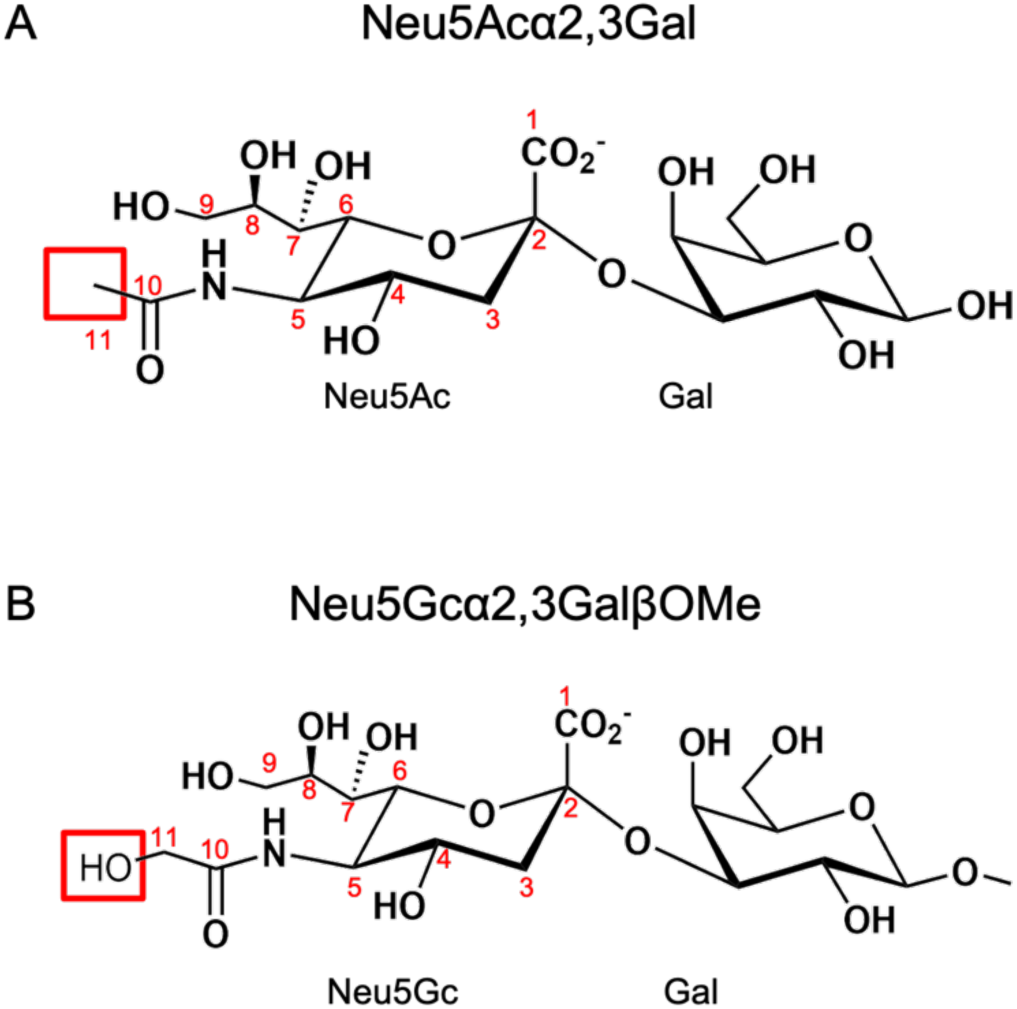
Chemical structures of Neu5Ac- and Neu5Gc-terminated α2-3-linked sialic acid-galactose disaccharides. **A)** α2-3-linked *N*-acetylneuraminic acid (Neu5Ac)-Galactose (Gal) (Neu5Acα2-3Gal). The C11 is highlighted. **B)** α2-3-linked *N*-glycolylneuraminic acid (Neu5Gc)-GalβOMe (Neu5Gcα2-3GalβOMe). The additional OH11 is highlighted.

Here, we evaluate SLBRs from *Streptococcus gordonii* strain Challis (SLBR_Hsa_) and *Streptococcus sanguinis* strain SK36 (SLBR_SrpA_), both of which can strongly engage sialoglycans capped by either Neu5Ac or Neu5Gc. Using structural and computational approaches, we asked how these SLBRs can bind synthetic sialoglycans terminating in either Neu5Ac or Neu5Gc, how this affects sialoglycan binding preference in solution, and how this correlates with native sialoglycoside engagement. Our results reveal that Neu5Ac versus Neu5Gc preference modulates, but does not uniquely determine, host specificity. These findings refine our understanding of how these bacteria target their hosts, clarify molecular aspects of tropism, and may even provide initial insight into molecular drivers of bacterial species jumps.

## Results

### X-ray Crystal Structures of SLBR_Hsa_ Bound to Neu5Ac- and Neu5Gc-Terminated Disaccharides

To investigate how SLBR_Hsa_ engages Neu5Ac versus Neu5Gc, we used X-ray crystallography. We determined X-ray crystal structures of SLBR_Hsa_ bound to synthetic α2-3-linked disaccharides Neu5Gcα2-3GalβOMe (SLBR_Hsa_–Neu5Gc, **Fig. 2A, 2B**) and Neu5Acα2-3Gal (SLBR_Hsa_–Neu5Ac, **Fig. 2A, 2C**). The SLBR_Hsa_–Neu5Gc complex was determined at 1.30 Å resolution (**Fig. 2B**, **Table 1**), and the SLBR_Hsa_–Neu5Ac complex was determined at 1.45 Å resolution (**Fig. 2C**, **Table 1**). Note that the methyl aglycon in Neu5Gcα2-3GalβOMe (**Fig. 2B**) arises from its chemical synthesis^30,31^. This leaving group is found on a region of the glycan that does not directly contact the SLBR (**Fig. 2B**). Moreover, we do not observe any structural perturbations attributable to this feature here or in reported costructures of Neu5Gc with SLBR_SrpA_^32^, where X-ray crystal structures with both Neu5Gc- and Neu5Ac-terminated α2-3-linked disaccharides have been reported^32,33^.

**Figure 2.**
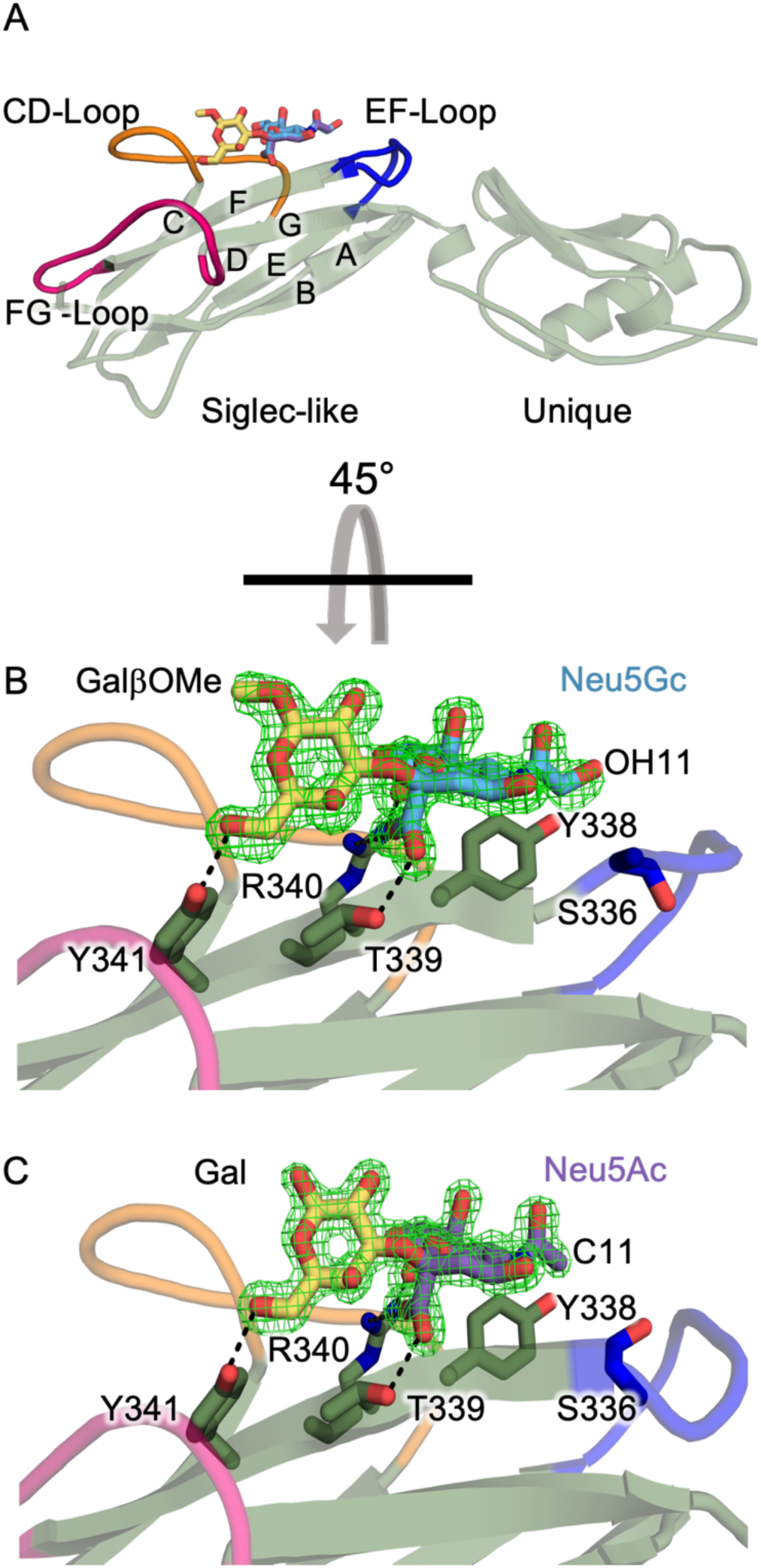
X-ray crystal structures of Neu5Acα2-3Gal or Neu5Gcα2-3GalβOMe bound to SLBR_Hsa_. **A)** Ribbons diagram of SLBR_Hsa_ with the strands of the V-set Ig fold labeled and the CD-(*orange*), EF-(*blue*), and FG-loops (*hot pink*) highlighted. Sialyl disaccharides are colored according to SNFG convention, with Neu5Ac in *purple*, Neu5Gc in *cyan*, and Gal in *yellow*. **B)** and **C)** Zoomed-in views rotated 45° around the x-axis to highlight hydrogen bonds between **B)** Neu5Acα2-3Gal or **C)** Neu5Gcα2-3GalβOMe and the ФTRX motif (SLBR_Hsa_^Y338^, SLBR_Hsa_^T339^, SLBR_Hsa_^R340^, SLBR_Hsa_^Y341^) on the F-strand. Each model is superimposed with composite omit electron density (*green mesh*).

**Table 1.** X-ray crystallographic data collection and refinement statistics for SLBR_Hsa_ bound to Neu5Acα2-3Gal or Neu5Gcα2-3GalβOMe. Values in parentheses are for the highest resolution shell. Raw data are deposited with SBGrid and can be accessed at: data.sbgrid.org/dataset/DATAID.

| Ligand | Neu5Acα2-3Gal | Neu5Gcα2-3GalβOMe |
| --- | --- | --- |
| PDB ID | 8ST5 | 8ST6 |
| DATA ID | 1020 | 1019 |
| Resolution | 1.45 Å | 1.30 Å |
| Highest resolution shell | (1.48 Å – 1.45 Å) | (1.32 Å – 1.30 Å) |
| *Data collection* |  |  |
| Beamline | SSRL 9-2 | SSRL 9-2 |
| Wavelength | 0.97946 Å | 0.97946 Å |
| Space group | P2<sub>1</sub>2<sub>1</sub>2<sub>1</sub> | P2<sub>1</sub>2<sub>1</sub>2<sub>1</sub> |
| Unit cell dimensions | a=46.1 Å, b=57.7 Å, c=76.0 Å | a=46.6 Å, b=58.1 Å, c=76.0 Å |
| R<sub>sym</sub> | 0.086 (0.490) | 0.079 (0.465) |
| R<sub>pim</sub> | 0.026 (0.204) | 0.023 (0.216) |
| I/$\sigma$ | 37.97 (1.81) | 48.1 (1.8) |
| Completeness (%) | 90.4% (47.1%) | 91.3% (46.5%) |
| Redundancy | 11.3 (5.9) | 11.0 (4.9) |
| CC<sub>1/2</sub> | 1.00 (0.964) | 1.00 (0.963) |
| *Refinement* |  |  |
| R<sub>cryst</sub> | 0.166 | 0.183 |
| R<sub>free</sub> | 0.198 (0.241) | 0.205 (0.288) |
| No. Mol per ASU | 1 | 1 |
| RMS deviation bond lengths | 0.010 | 0.004 |
| RMS deviation bond angles | 0.995 | 0.681 |
| Ramachandran Statistics |  |  |
| Favored | 95.61% | 97.54% |
| Outliers | 0.49% | 0.00% |

The positions and binding of these two sialyl disaccharide ligands are conserved between SLBR_Hsa_–Neu5Ac and SLBR_Hsa_–Neu5Gc, with superposition of the SLBR_Hsa_–Neu5Ac and SLBR_Hsa_–Neu5Gc showing that the binding position for each sialoglycan is within the error of the resolution. Each sialoglycan binds above the conserved ΦTRX motif of the F strand of the Siglec domain, where residues SLBR_Hsa_^Y338^, SLBR_Hsa_^T339^, SLBR_Hsa_^R340^, and SLBR_Hsa_^Y341^ stabilize ligands with hydrogen-bonding interactions (**Fig. 2B, 2C**). This binding site is located between the CD, EF, and FG loops, making the overall binding mode similar to the Neu5Acα2-3Gal terminus of the sialyl tri- and tetra-saccharides in reported SLBR–sialoglycan complexes^2,16^ (**Fig. 2B, 2C**).

Structural comparison of SLBR_Hsa_–Neu5Ac and SLBR_Hsa_–Neu5Gc with the previously reported unliganded SLBR_Hsa_^16^ also reveals similar overall folds, with RMSD values for Cα atoms of 0.157 Å (Neu5Ac-bound) and 0.158 Å (Neu5Gc-bound). One small difference is the position of the EF loop. Past work shows that this EF loop can close over bound tri- and tetrasaccharides^16^. In these structures, the EF loop closes over Neu5Gc but not over Neu5Ac, with a maximal backbone displacement of 4.6 Å (**Fig. 2B, 2C, Supplementary Fig. 1A**). Loop closure is unlikely to impact Neu5Ac versus Neu5Gc selectivity as it does not detectably affect contacts to C11 or the C11-appended hydroxyl (OH11). Moreover, past work using the same crystal form of SLBR_Hsa_ identified that the EF loop is somewhat stabilized in the open position by crystal contacts^16^. The difference in loop position that is observed here more likely results from this same phenomenon rather than by differences in ligand.

To provide more insight, we compared these SLBR_Hsa_ structures to reported structures of SLBR_SrpA_, a related SLBR that can similarly bind sialoglycans terminating in either Neu5Ac or Neu5Gc^7,33^ and where experimental structures with each ligand have been reported^32^ (**Fig. 3, Supplementary Fig. 1B, Supplementary Fig. 2**). Comparison of SLBR_Hsa_–Neu5Gc with SLBR_SrpA_–Neu5Gc^32^ shows that Neu5Gc similarly binds above the F strand in both these SLBRs, albeit with a lateral shift in position of 1.5 Å with respect to the ΦTRX motif (**Supplementary Fig. 1B**). An additional difference is a 160° rotation of OH11 between the two ligands (**Fig. 3A, 3B**). In SLBR_Hsa_–Neu5Gc, the OH11 orients toward solvent and does not interact with the protein (**Fig. 3A**). The closest protein atoms to OH11 are the SLBR_Hsa_^Y338^ side chain hydroxyl on the F-strand and SLBR_Hsa_^S336^ Oγ on the EF loop, with distances of 3.8 Å in each case (**Fig. 3A**). The closest atoms to C11 are the SLBR_Hsa_^Y338^ side chain hydroxyl and SLBR_Hsa_^S336^ Cβ, with distances of 3.2 Å and 3.9 Å, respectively (**Fig. 3A**). In comparison, a unique hydrogen bond forms between OH11 and the SLBR_SrpA_^Y368^ side chain hydroxyl that orients OH11 toward the protein (**Fig. 3B**). Comparison of SLBR_Hsa_–Neu5Ac (**Supplementary Fig. 2A**) with SLBR_SrpA_–Neu5Ac^32^ (**Supplementary Fig. 2B**) shows the same 1.5 Å lateral shift in glycan position with respect to the ΦTRX motif as is observed for the Neu5Gc costructures.

**Figure 3.**
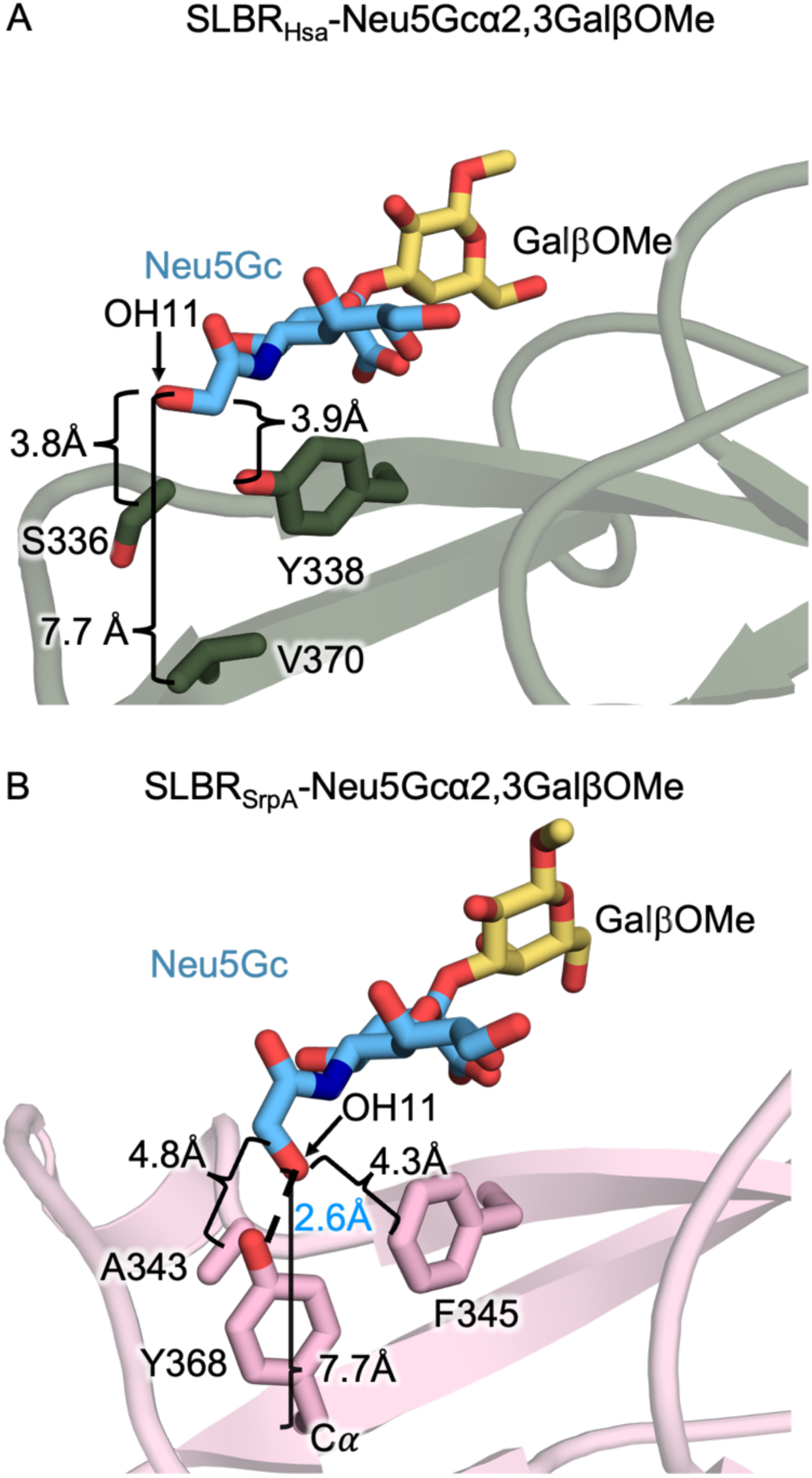
Structures of SLBR_Hsa_–Neu5Gc and SLBR_SrpA_–Neu5Gc. Side-by-side comparison of Neu5Gc-GalβOMe binding in **A)** SLBR_Hsa_ and **B)** SLBR_SrpA_^32^ illustrates differences in the positioning of the C11 hydroxyl group (OH11) of Neu5Gc. **A**) In SLBR_Hsa_, the residues adjacent to sialic acid include SLBR_Hsa_^S336^ and SLBR_Hsa_^Y338^. These residues lack direct contact with the OH11 group, which is oriented away from the binding site. **B**) In contrast, SLBR_SrpA_^Y368^ hydrogen-bonds to the OH11 group. Distances for adjacent non-bonding atoms are indicated with brackets.; the hydrogen-bond between SLBR_SrpA_^Y368^ OH and Neu5Gc OH11 is shown with a dotted line and marked with blue text.

We next assessed whether there were differences between the Neu5Ac/Neu5Gc binding SLBRs and Neu5Ac-selective SLBRs (**Fig. 4, Supplementary Fig. 2**). For the Neu5Ac/Neu5Gc binding SLBRS, we used SLBR_Hsa_–Neu5Ac/Neu5Gc (**Fig. 4A, Supplementary Fig. 2A**), SLBR_SrpA_–Neu5Ac/Neu5Gc (**Fig. 4B, Supplementary Fig. 2B**) and an additional Neu5Ac/Neu5Gc binding comparator, SLBR_UB10712_ (**Fig. 4C**) where there is only a structure of the SLBR without ligand bound^16^. For the Neu5Ac-selective SLBRs, we used SLBR_GspB_–Neu5Ac^14^ (**Fig. 4D, Supplementary Fig. 2C**), SLBR_SK1_–Neu5Ac^34^ (**Fig. 4E, Supplementary Fig. 2D**), and SLBR_SK678_^16^ (**Fig. 4F**), which only has a structure of the SLBR without ligand bound^16^. This comparison identified that the Neu5Ac/Neu5Gc-binding SLBRs had a more open binding pocket near C11 and OH11 (**Fig. 4A, 4C, 4E**) while the Neu5Ac-selective SLBRS shows a more defined pocket that likely has more precise Van der Waals contacts near C11 (**Fig. 4B, 4D, 4F**). Other potential features such as electrostatics (**Fig. 4A-4F**), bonding patterns, and water molecule substructure did not correlate with sialic acid selectivity.

**Figure 4.**
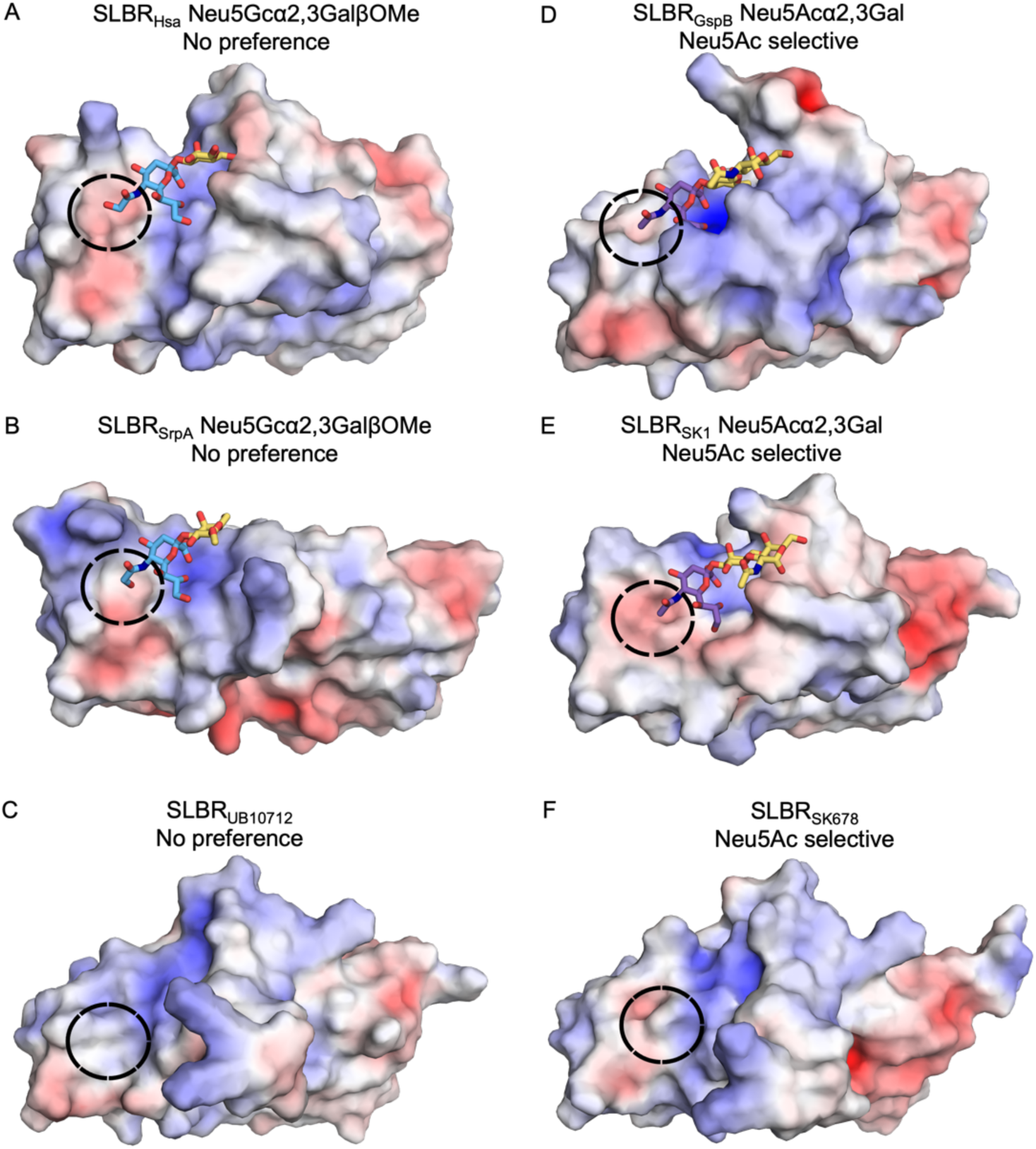
Surface rendering of SLBRs colored by electrostatic potential. In the Figure, surfaces with positive charge are colored *blue*, surfaces with negative charge are colored *red*, and neutral surfaces are colored *white*. **A)** SLBR_Hsa_, **B)** SLBR_SrpA_^32^, **C)** SLBR_UB19712_^16^, **D)** SLBR_GspB_^14^, **E)** SLBR_SK1_^34^, and **F)** SLBR_SK678_^16^. The black circle highlights the surface adjacent to C11/OH11 and shows a more defined binding pocket in the characterized Neu5Ac-selective SLBRs.

### Molecular Dynamics (MD) Simulations of SLBR_Hsa_ and Glycan Motions

To calculate whether motions in SLBR_Hsa_ might affect interactions with Neu5Ac- or Neu5Gc-terminated sialoglycans, we conducted molecular dynamics (MD) simulations. In these simulations, we separately calculated SLBR_Hsa_ flexibility and sialyl disaccharide flexibility. The coordinates for Neu5Gcα2-3Gal did not include the methyl aglycon, so both sialoglycans had equivalent structures.

Simulations of SLBR_Hsa_ flexibility were initiated from the unliganded conformation of SLBR_Hsa_^16^, allowing the loops to equilibrate around each ligand without pre-imposed bias. As in prior simulations^16^, the CD-, EF-, and FG-loops adjacent to the binding pocket exhibited the greatest flexibility, supported by both root-mean-square fluctuation (RMSF) (**Supplementary Fig. 3**) and crystallographic temperature factor analyses (**Supplementary Fig. 4**). In addition, the EF loop of SLBR_Hsa_ closed over both Neu5Acα2-3Gal and Neu5Gcα2-3Gal, allowing the SLBR_Hsa_^K335^ backbone carbonyl to approach the O4 atom of each sialoglycan (**Supplementary Fig. 5**). This differs from what was observed in the X-ray crystal structures, in which only the SLBR_Hsa_–Neu5Gc structure had a closed loop (**Supplementary Fig. 1A**). We particularly evaluated SLBR_Hsa_^V370^ on the G strand, which is analogous to the SLBR_SrpA_^Y368^ that hydrogen-bonds with Neu5Gc OH11. SLBR_Hsa_^V370^ did not exhibit flexibility or approach either sialoglycan. In fact, no protein atoms came within 3 Å of the Neu5Ac/Neu5Gc C11 or Neu5Gc OH11 at any point during the simulation. These predictions suggest that, on the timescale of the simulation, motions in SLBR_Hsa_ do not promote stable direct contacts with C11 or OH11.

We designed a second set of calculations to evaluate whether sialoglycan flexibility might contribute to transient interactions with SLBR_Hsa_ by allowing C11, OH11, or other atoms to approach the protein surface (**Supplementary Fig. 6**). These simulations were initiated with the EF loop of SLBR_Hsa_ in the closed conformation around each disaccharide. Throughout the simulations, the Neu5Acα2-3Gal and Neu5Gcα2-3Gal disaccharides remained stably bound with a conserved orientation. The sialic acid and galactose moieties showed minimal positional fluctuation. In the Neu5Gc-bound simulation, the OH11 did not form persistent contacts with the protein surface. These findings are consistent with the crystal structure of SLBR_Hsa_–Neu5Gc, in which OH11 is rotated away from the protein and SLBR_Hsa_ does not directly engage Neu5Gc OH11 (**Fig. 3**). The resulting probability distributions in the replicates showed show a narrow probability distribution around the pose of the crystal structure (**Supplementary Fig. 6**). The mean RMSD is 0.85 Å ± 0.14 Å for Neu5Ac and 0.83 Å ± 0.15 Å for Neu5Gc.

### Binding of SLBR_Hsa_^V170^ and SLBR_SrpA_^Y368^ Mutants to Purified Glycans Synthetic Disaccharides

We next explored the potential for indirect influences that might help SLBRs distinguish between these two forms of sialic acid. We focused on residues analogous to SLBR_SrpA_^Y368^, whose side chain hydroxyl forms a hydrogen bond with the OH11 group of Neu5Gc (**Fig. 3B**). The Cα of SLBR_SrpA_^Y368^ is 7.6 Å from OH11 and the close approach of the side chain hydroxyl is due to the length of the residue (**Fig. 3B**). SLBR_SrpA_^Y368^ corresponds with SLBR_Hsa_^V370^ (**Fig. 3A, 3B**). While SLBR_SrpA_^Y368^ makes a hydrogen bond to the Neu5Gc OH11 (**Fig. 3B**), other amino acids at this position are not capable of a hydrogen-bonding interaction. Among characterized SLBRs, residues at the equivalent position are typically hydrophobic: Val, Ile, Tyr, or Phe (**Fig. 5**). These residues are also smaller than Tyr. For example, the closest side chain atoms from the shorter SLBR_Hsa_^V370^ 7.5 Å away from C11 (**Fig. 3A**, **Supplementary Fig. 2A**). Finally, except for SLBR_SrpA_^Y368^, these analogous residues do not contact the ligand. For example, SLBR_Hsa_^S336^ and SLBR_Hsa_^Y338^ are located between the SLBR_Hsa_^V370^ side chain and C11, blocking the possibility of direct contact (**Fig. 3A**).

**Figure 5.**
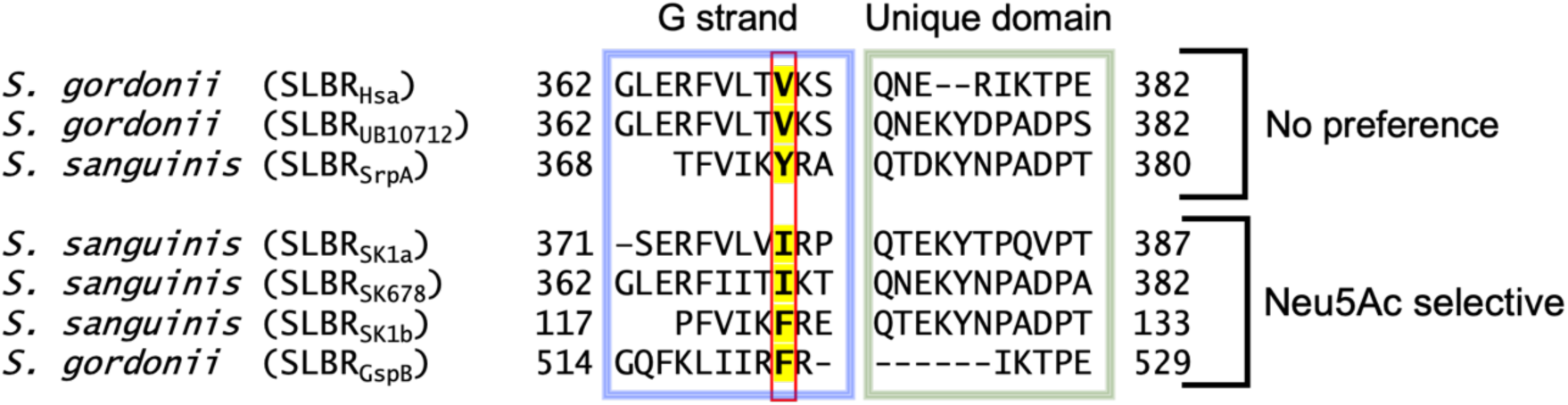
Structure-based sequence alignment of SLBRs with reported Neu5Ac/Neu5Gc binding data. Sequences are from WP_081102781.1 from *S. gordonii* strain Challis (SLBR_Hsa_)^16,61^, WP_045635027.1 from *S. gordonii* strain UB10712^16,62^, WP_011836739.1 from *S. sanguinis* strain SK36 (SLBR_SrpA_)^32,63^, WP_080555651.1 from *S. sanguinis* strain SK1 (SLBR_SK1a_ and SLBR_SK1b_)^34,64^, WP_125444035.1 from *S. sanguinis* strain SK678^16,64^, and WP_125444382.1 from *S. gordonii* strain M99 (SLBR_GspB_)^14,65^. The highlighted residue is mutated in this study.

To investigate whether residues at this position play an allosteric role in distinguishing between sialic acid forms, we made substitutions in SLBR_Hsa_ and SLBR_SrpA_. SLBR_Hsa_^V370^ was mutated to Ile, Phe, or Tyr and SLBR_SrpA_^Y368^ was mutated to Val, Ile, or Phe, so that each of these SLBR scaffolds had a version with Val, Ile, Tyr, or Phe at this position. We assessed binding to purified and biotinylated Neu5Acα2-3Gal or Neu5Gcα2-3Gal by ELISA (**Fig. 6, 7**), evaluating both the overall level of binding and the preference for Neu5Ac-versus Neu5Gc-terminated disaccharides. Consistent with previous reports, wild-type SLBR_Hsa_ bound sialyl disaccharides at much higher levels than SLBR_SrpA_, but each wild-type SLBR had no preference for sialyl disaccharides terminating in Neu5Ac or Neu5Gc (**Fig. 6, 7**).

**Figure 6.**
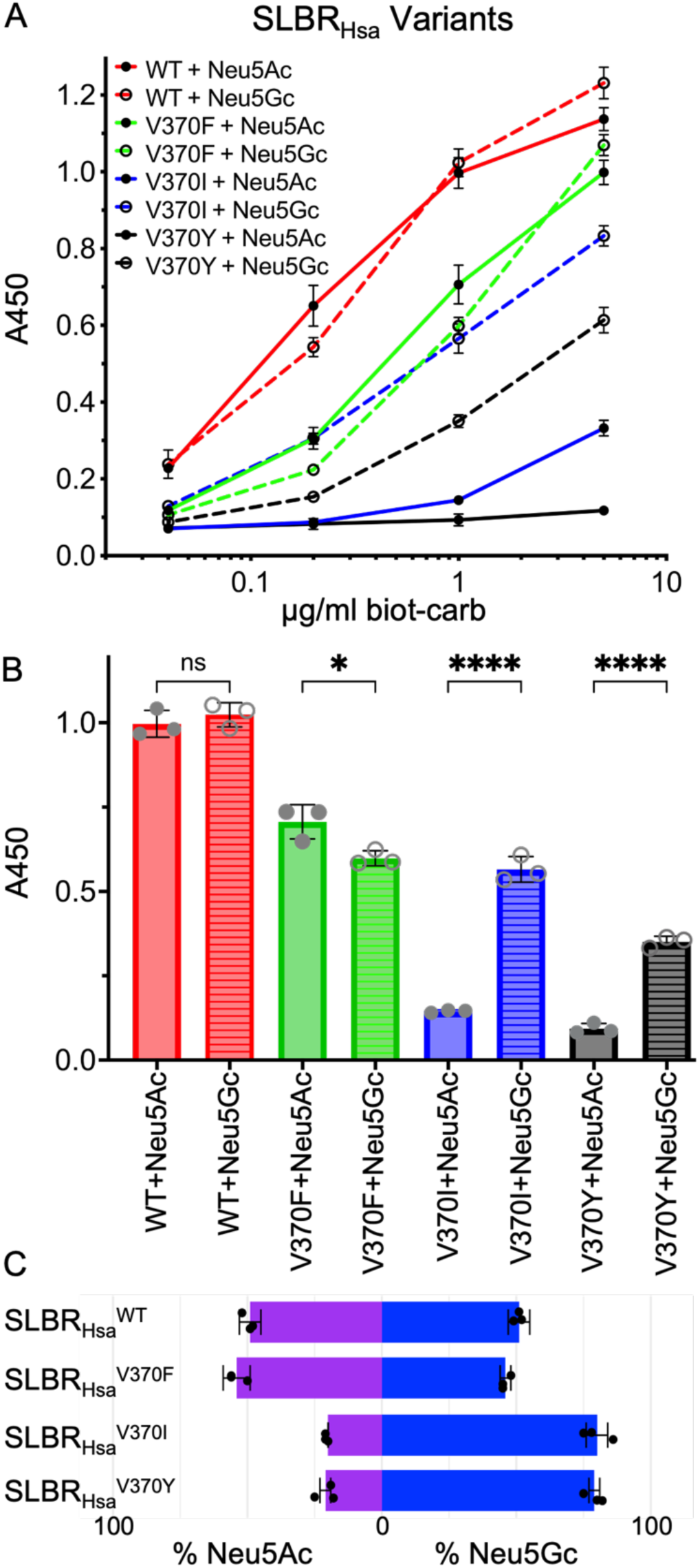
Wildtype and mutant SLBR_Hsa_ binding to purified biotinylated disaccharides. **A)** ELISA curves for wildtype and mutant GST-tagged SLBR_Hsa_ (GST-SLBR_Hsa_) binding to biotinylated Neu5Ac- and Neu5Gc-terminated disaccharides at the indicated concentrations. Measurements were performed using 500 nM of immobilized GST-SLBR_Hsa_, and the indicated concentrations of each ligand are shown as the mean ± SD. (n = 3 independent experiments performed on protein from a single preparation). **B)** Comparison of the levels of binding of wildtype and mutant GST-SLBR_Hsa_ at a concentration of 1 μg/mL biotinylated disaccharides. p-values for Neu5Ac versus Neu5Gc binding to SLBR_Hsa_ are: SLBR_Hsa_^WT^ p= 0.9585, SLBR_Hsa_^V370F^ p= 0.0121, SLBR_Hsa_^V370I^ p= <0.0001, SLBR_Hsa_^V370Y^ p= <0.0001, as evaluated by one-way ANOVA, **C)** Preference of wildtype or mutant GST-SLBR_Hsa_ for Neu5Ac (*purple*) versus Neu5Gc (*blue*). Percentages were calculated by dividing measured A_450_ values corresponding to either Neu5Ac or Neu5Gc by the additive absorbance value (Neu5Ac+Neu5Gc).

As compared to wild-type SLBR_Hsa_, SLBR_Hsa_^V370I^ and SLBR_Hsa_^V370Y^ exhibited reduced but statistically significant Neu5Acα2-3Gal binding, while SLBR_Hsa_^V370Y^ lacked statistically significant binding under the conditions tested (**Fig. 6A, 6B**). In contrast, all SLBR_Hsa_ proteins retained binding to Neu5Gcα2-3Gal, although overall levels were moderately reduced (**Fig. 6A, 6B**). Because of the disproportionate loss of Neu5Acα2-3Gal binding, all SLBR_Hsa_ mutants became more selective for Neu5Gc-terminated disaccharides (**Fig. 6C**).

Mutations in SLBR_SrpA_ produced an even more striking result. All variants showed substantially increased relative binding to Neu5Gc-termined disaccharides when compared to the wild-type SLBR_SrpA_. The strongest effect was observed with the SLBR_SrpA_^Y368I^, where binding to Neu5Gcα2-3Gal increased 5.1-fold when 1 µg/mL of each biotinylated disaccharide was used (**Fig. 7A, 7B**). The SLBR_SrpA_^Y368V^ (3.9-fold increase) and SLBR_SrpA_^Y368F^ (3.9-fold increase) mutants also showed substantial increases in Neu5Gcα2-3Gal binding (**Fig. 7A, 7B**). At the same time, Neu5Acα2-3Gal binding increased 2.7-fold in SLBR_SrpA_^Y368F^ but was statistically identical in the remaining mutants. Because of the very large gains in Neu5Gcα2-3Gal binding, SLBR_SrpA_ mutants were even more strongly selective for Neu5Gc-terminated disaccharides than were SLBR_Hsa_ mutants (**Fig. 7C**).

**Figure 7.**
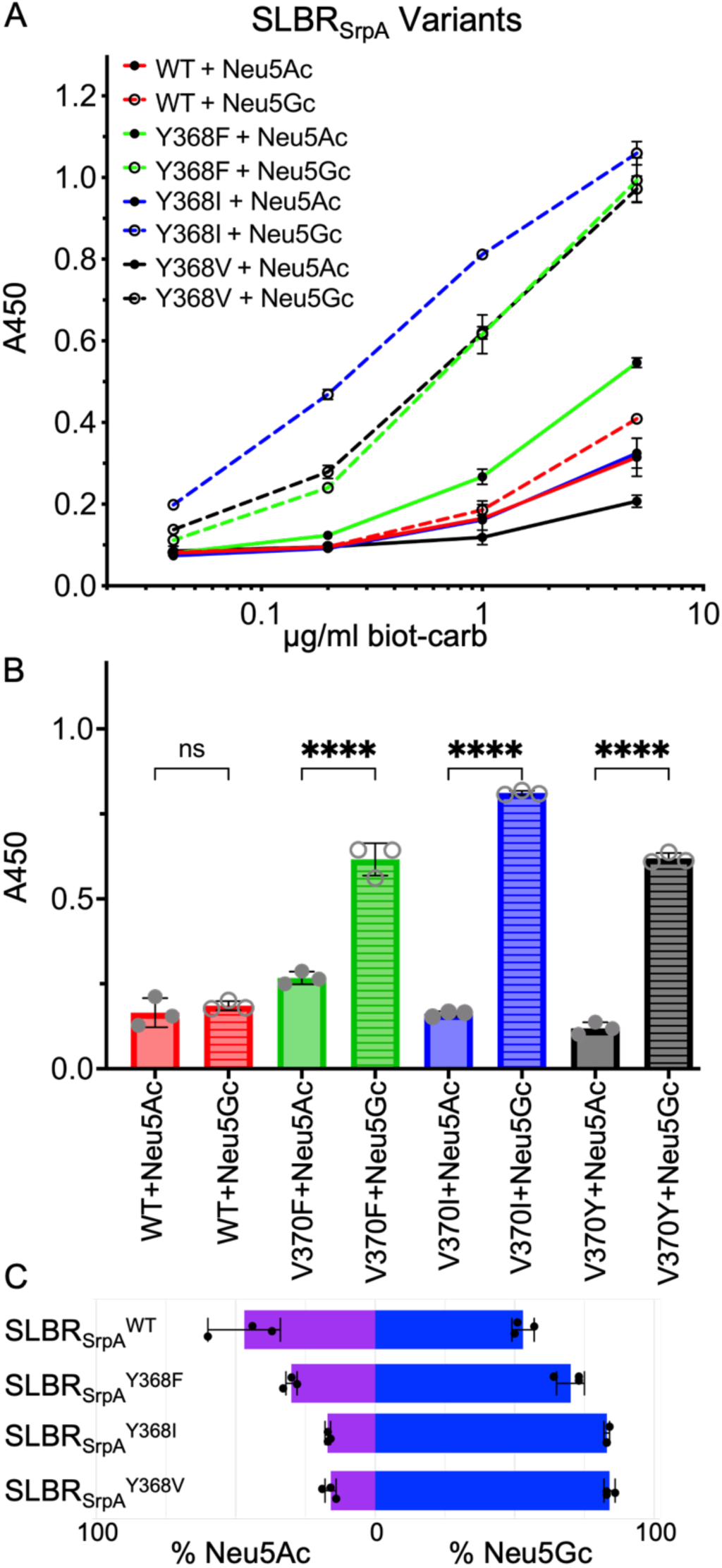
Wildtype and mutant SLBR_SrpA_ binding to purified and biotinylated disaccharides. **A)** ELISA curves for wildtype and mutant GST-tagged SLBR_Hsa_ (GST-SLBR_Hsa_) binding to biotinylated Neu5Ac- and Neu5Gc-terminated disaccharides at the indicated concentrations. Measurements were performed using 500 nM of immobilized GST-SLBR, and the indicated concentrations of each ligand are shown as the mean ± SD. (n = 3 independent experiments performed on protein from a single preparation). **B)** Comparison of the levels of wildtype and mutant GST-SLBR_SrpA_ binding to 1 μg/mL biotinylated Neu5Ac versus Neu5Gc disaccharides. p-values for Neu5Ac versus Neu5Gc binding to SLBR_SrpA_ are: SLBR_SrpA_^WT^ p= 0.9407, SLBR_SrpA_^Y368F^ p= <0.0001, SLBR_SrpA_^Y368I^ p= <0.0001, SLBR_SrpA_^Y368V^ p= <0.0001, as evaluated by one-way ANOVA, **C)** Preference of wildtype or mutant GST-SLBR_SrpA_ for Neu5Ac (*purple*) versus Neu5Gc (*blue*). Percentages were calculated by dividing measured A_450_ values corresponding to either Neu5Ac or Neu5Gc by the additive absorbance value (Neu5Ac+Neu5Gc).

Taken together, our mutational analysis identified that the SLBR_Hsa_^V370^ and SLBR_SrpA_^Y368^ positions affect sialic acid selectivity and that all tested mutations at this position increased Neu5Gc selectivity. The substantial gains in Neu5Gc preference were particularly intriguing because the substitutions maintain hydrophobicity near the sialoglycan binding site, and Neu5Gc is more hydrophilic.

### Far Western Analysis with Wild-type and Neu5Gc-selective mutants

We next investigated binding to authentic glycoproteins from human and rat sources by far Western analysis (**Fig. 8, 9**). Within the limits of detection, human plasma contains glycoproteins that only terminate in Neu5Ac and its derivatives, while rats have glycoproteins terminating in both Neu5Ac and Neu5Gc^23^. In addition, this evaluates how selectivity for Neu5Ac versus Neu5Gc is combined with potential contributions from glycan branching or protein context.

**Figure 8.**
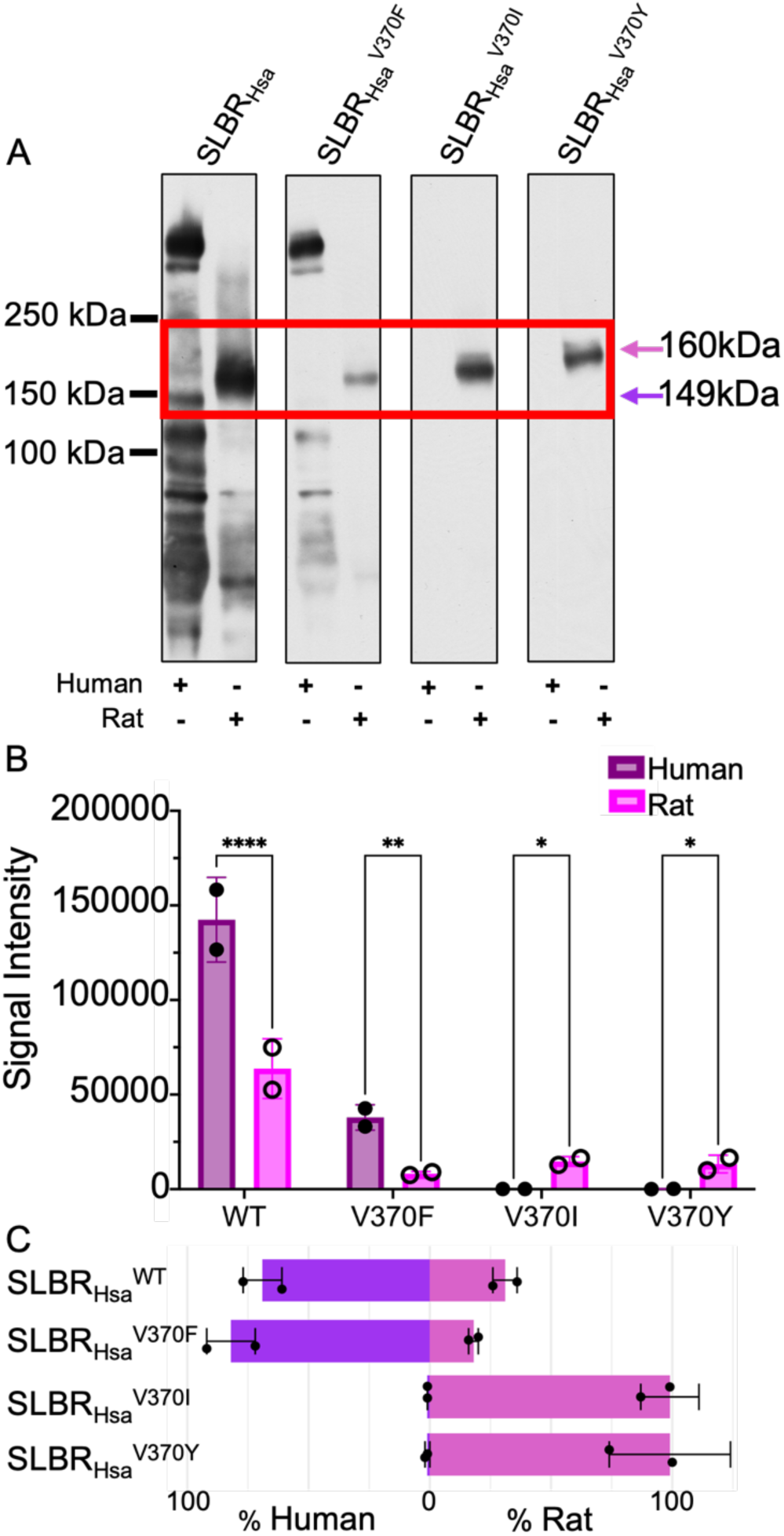
SLBR_Hsa_ binding to human or rat plasma glycoproteins. **A)** Far-Western blot of wild-type and mutant GST-SLBR_Hsa_ against plasma glycoproteins. Glycoproteins were separated by electrophoresis through a 3–8% polyacrylamide gradient, and then stained. No signals were detected outside of the cropped region. As previously identified^6^, the proteins highlighted by the red box are human GPIbα (149 kDa, *purple arrow*) or rat GPIbα (160 kDa, *pink arrow*). **B)** Dosimetry of blots showing the total amount of binding to all proteins, without regard to molecular weight, measured with IMAGEJ version 1.54^66^. p-values for human versus rat binding to SLBR_Hsa_ are: SLBR_Hsa_^WT^ p= <0.0001, SLBR_Hsa_^V370F^ p= 0.0012, SLBR_Hsa_^V370I^ p= 0.0155, SLBR_Hsa_^V370Y^ p= 0.0214, as evaluated by two-way ANOVA, **C)** Preference of wildtype or mutant GST-SLBR_Hsa_ for human (Neu5Ac only, *purple*) versus rat (Neu5Ac/Neu5Gc, *pink*). Percentages were calculated by dividing pixel counts corresponding to either human or rat samples by the additive pixel counts (human + rat).

Contrary to what was observed in ELISAs with synthetic sialoglycans (**Fig. 6**), wild-type SLBR_Hsa_ showed a robust interaction with the human plasma glycoproteins (**Fig. 8A, 8B**) and a substantial preference for binding to glycoproteins within human plasma (**Fig. 8C**). However, there are some curious nuances. Of particular note is binding to the ∼150 kDa GPIbα glycoprotein implicated as the receptor in endocardial infection (**Fig. 8A**). SLBR_Hsa_ exhibits more total binding to human plasma glycoproteins than rat plasma glycoproteins (**Fig. 8B**), but much of this is off target with respect to GPIbα (**Fig. 8A**). While there is less total binding of SLBR_Hsa_ to rat plasma glycoproteins, this SLBR more selectively recognizes rat GPIbα (**Fig. 8A**).

Among SLBR_Hsa_ mutants, SLBR_Hsa_^V370F^ retained a preference for human plasma glycoproteins, albeit at a 3-fold decrease in total binding (**Fig. 8A, 8B**). SLBR_Hsa_^V370F^ also lost detectable binding to human GPIbα and only bound to off-target glycoproteins not associated with endocarditis (**Fig. 8A**). All detectable binding of SLBR_Hsa_^V370F^ to rat plasma glycoproteins is to GPIbα (**Fig. 8A**). The remaining two SLBR_Hsa_ mutants only detectably bound to rat GPIbα (**Fig. 8A**).

Wild-type SLBR_SrpA_ bound more robustly to rat plasma (**Fig. 9A, 9B**), again in contrast to the statistically identical binding to Neu5Ac- and Neu5Gc-terminated synthetic disaccharides in ELISA (**Fig. 7**). Moreover, almost all detectable SLBR_SrpA_ binding was to GPIbα in both human and rat plasma (**Fig. 9A**). Of the SLBR_SrpA_ mutants, SLBR_SrpA_^Y368V^ and SLBR_SrpA_^Y368I^ either had statistically identical or modestly increased the binding to binding to rat GPIbα (**Fig. 9A, 9B**), which contrasted with the large increase of binding of each of these SLBR mutants to purified Neu5Gc-terminated disaccharides (**Fig. 7**). For example, wild-type SLBR_SrpA_ and SLBR_SrpA_^Y368F^ had statistically identical binding to rat GPIbα (**Fig. 9A, 9B**), even though SLBR_SrpA_^Y368F^ had 5-fold greater binding to synthetic Neu5Gc-terminated disaccharides in ELISA (**Fig. 7A, 7B**). The SLBR_SrpA_^Y368F^ mutant showed barely detectable binding to human plasma glycoproteins, despite a ∼3-fold increase in binding to Neu5Acα2-3Gal binding in the ELISA assays (**Fig. 9**). The remaining two SLBR_SrpA_ mutants had undetectable binding to human plasma glycoproteins under these conditions, and all SLBR_SrpA_ mutants substantially shifted the binding preference toward rat glycoproteins (**Fig. 9C**).

**Figure 9.**
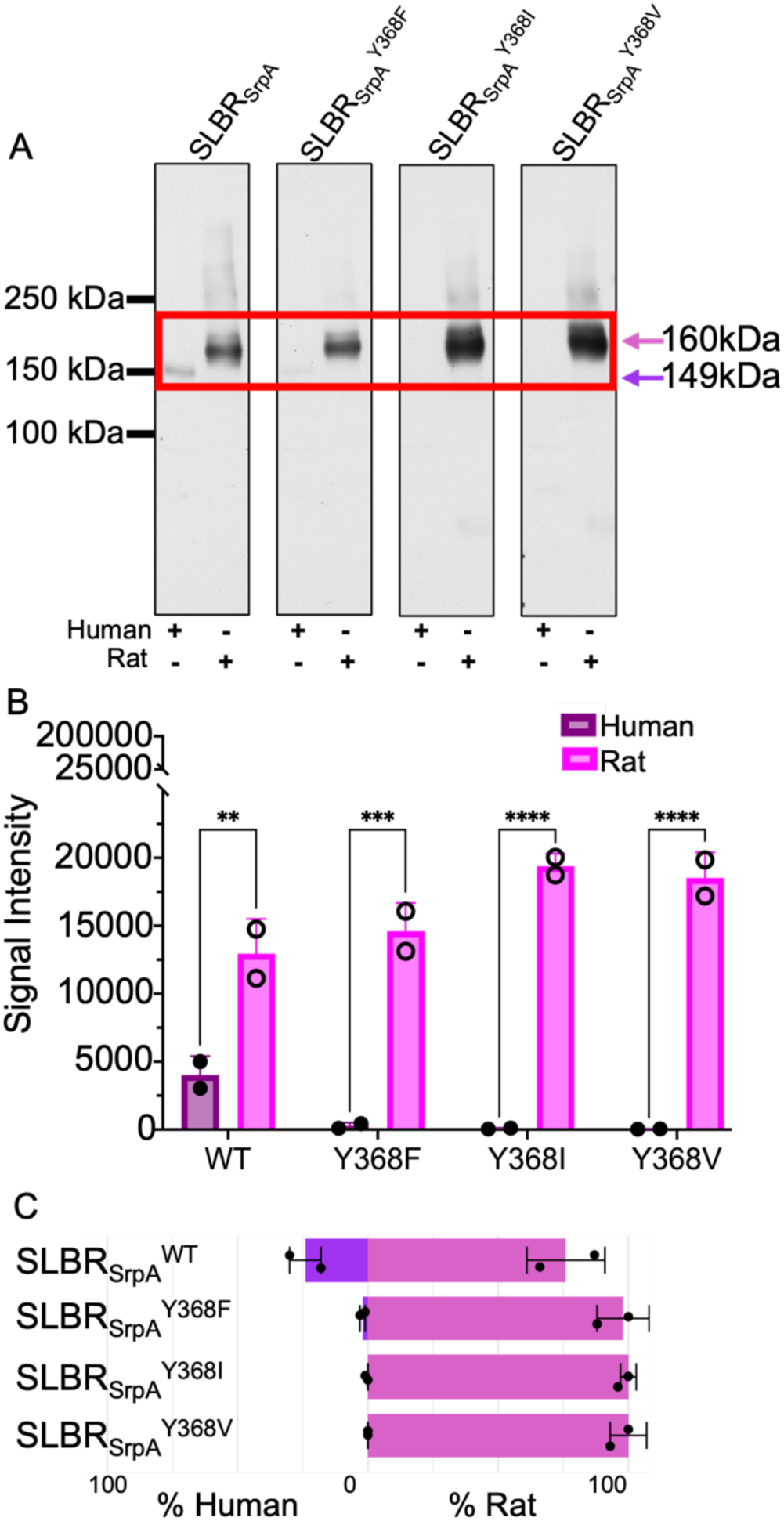
Far Western analysis of SLBR_SrpA_ binding to human or rat plasma. **A)** Far-Western blot of wild-type and mutant GST-SLBR_SrpA_ against plasma glycoproteins. Glycoproteins were separated by electrophoresis through a 3–8% polyacrylamide gradient, and then stained. No signals were detected outside of the cropped region. The proteins in the red box are human GPIbα (149 kDa, *purple arrows*) or rat (160 kDa, *pink arrows*) GPIbα^6^. **B)** Dosimetry of blots showing the total amount of binding to all proteins, without regard to molecular weight, measured with IMAGEJ version 1.54^66^. p-values for human versus rat binding to SLBR_SrpA_ are: SLBR_SrpA_^WT^ p= 0.0011, SLBR_SrpA_^Y368F^ p= 0.0002, SLBR_SrpA_^Y368I^ p= <0.0001, SLBR_SrpA_^Y368V^ p= <0.0001, as evaluated by two-way ANOVA, **C)** Preference of wildtype or mutant GST-SLBR_Hsa_ for human (Neu5Ac only, *purple*) versus rat (Neu5Ac/Neu5Gc, *pink*). Percentages were calculated by dividing pixel counts corresponding to either human or rat samples by the additive pixel counts (human + rat).

## Discussion

Our data provide insight into how SLBRs from viridans group streptococci engage Neu5Ac- and Neu5Gc-terminated sialoglycans (**Fig. 1**). We define the structural basis for binding of SLBR_Hsa_ to the Neu5Gcα2-3Gal and Neu5Gcα2-3Gal sialyl disaccharides (**Fig. 2-4, Supplementary** Fig 1**, 2**), show that motions are unlikely to affect Neu5Ac/Neu5Gc selectivity (**Supplementary** Fig 3-6), and demonstrate that Neu5Ac/Neu5Gc preference can be modified through mutation of an allosteric site (**Fig. 5-7**). Furthermore, we show that that binding to synthetic Neu5Ac- and Neu5Gc-terminated disaccharides only partially predicts binding to authentic human and rat plasma sialoglycoproteins (**Fig. 6-9**). Intriguingly, SLBR mutants also showed narrowed glycoprotein engagement (**Fig. 8, 9**), losing off-target binding and very selectively engaging the GPIbα glycoprotein that is implicated as the receptor for infective endocarditis^7,35,36^. Several aspects of these results suggest important nuances in how SLBRs mediate host glycoprotein recognition.

A key finding was that Neu5Ac/Neu5Gc selectivity appears to be modulated by allosteric effects rather than by direct contacts (**Fig. 2, 3**). In addition, there was no evidence that flexibility contributes meaningfully to Neu5Ac/Neu5Gc cross-reactivity (**Supplementary Fig. 3-6**). This fully contrasts with how SLBRs distinguish between tri- and tetrasaccharide sialoglycans with different chemical compositions, in which binding preferences are largely dictated by direct SLBR-ligand interactions, as assisted by protein flexibility^16^. In those cases, chimeragenesis and point mutagenesis resulted in predictable changes to the sialoglycan binding repertoire^16^ and flexibility of the CD-, EF-, and FG-loops (**Fig. 2A**) were shown to contribute to broad selectivity^16^.

Another striking finding was the consistent gain in Neu5Gc preference for both SLBRs following point mutagenesis (**Fig. 6-9**). This asymmetry may arise from differences in the requirements for each form of sialic to interact with SLBRs (**Fig. 2-4**). A polar OH11 of Neu5Gc may benefit from a more open binding pocket (**Fig. 4A-4C**) to allow local motions and enhanced solvent exposure. This may be recapitulated by perturbed local packing that inevitably accompanies mutagenesis. A hydrophobic Neu5Ac C11 methyl group may require more precise packing and desolvation (**Fig. 4D-4F**). Despite this, all naturally occurring SLBRs that have been experimentally characterized either exhibit a strong binding preference toward sialoglycans terminating in Neu5Ac^2,16,19^ or bind equivalently to sialoglycans terminating in Neu5Ac or Neu5Gc^2^ (**Fig. 6A, 7A**). While it is not clear why this is, one possibility is that the absence of characterized SLBRs that prefer Neu5Gc-terminated sialoglycans may reflect that the library of available viridans group streptococci favors bacteria isolated from humans^2–4^. These isolates face selective pressure to engage Neu5Ac-terminated sialoglycosides of humans^5,6^.

The imperfect correlation between Neu5Ac/Neu5Gc selectivity (**Fig. 6,7**) and host preference (**Fig. 8,9**) extends our understanding of host tropism by sialoglycan-binding pathogens. One hypothesis for the divergence in sialic acid composition between humans and non-human animals involves the evolutionary response to host-pathogen interplay. Non-human animals synthesize CMP-linked Neu5Gc when CMP-Neu5Ac hydroxylase enzyme hydroxylates the *N*-acetyl group (i.e. C11) of CMP-Neu5Ac. Our study used samples from rats, where Neu5Gc levels have been experimentally measured at different levels depending on the biological location. The ratio in developing lungs measured as ∼75% Neu5Ac and ∼25% Neu5Gc^23^ while the level in rat GPIbα extract is measured as ∼28% Neu5Ac and ∼58% Neu5Gc^6^. Humans have an inactive CMP-Neu5Ac hydroxylase enzyme due to a loss-of-function mutation occurring approximately two million years ago^21,25,37,38^, and therefore only synthesize Neu5Ac. It has been postulated that this human loss of function mutation conferred protection against ancestral forms of *Plasmodium*, the parasitic protozoan responsible for malaria^39^ as some modern *P. falciparum* strains rely primarily on sialic acid for invasion^40,41^.

These results support a model where the relationship between pathogenesis and sialic acid identity is not straightforward. Instead linkage specificity (e.g., α2-3 vs α2-6 of C6 to the underlying Gal) may play a large role than sialic acid identity in determining virulence. This aligns well with other characterized pathogens that bind to sialoglycans. One example is observed in the influenza virus, which binds to host sialoglycans through hemagglutinins. The H1 and H3 hemagglutinins specifically bind α2-6 linked sialic acids, which are abundant in mammalian airways^42^. These strains are infectious to humans and other mammals. By contrast, H5 hemagglutinin found in strains of the avian flu binds to α2-3 linked sialic acids, which are abundant in avian airways but rare in mammalian airways^43,44^. The H5N1 avian flu has a high mortality rate in both mammals and birds, but H5N1 strains are substantially less infective for humans at the present time because of this linkage selectivity^45^. Intriguingly, Neu5Gc-terminated sialoglycans can act as decoy receptors for some strains of influenza, where they support hemagglutinin binding but inhibit viral entry^35–37^. Coronaviruses such as SARS-CoV and SARS-CoV2 can also engage host sialoglycans^46–48^, and this interaction synergizes with binding to ACE2 to promote viral entry^46–48^. Although this finding is quite recent, early work similarly suggests narrow linkage selectivity for this interaction^46–48^.

Our results suggest that host tropism of viridans group streptococci blends Neu5Ac/Neu5Gc selectivity with broader glycan context. Host glycoprotein engagement is certainly influenced by the ability to bind Neu5Ac versus Neu5Gc-capped sialoglycans, as evidenced by shifts in SLBR binding toward rat glycoproteins as Neu5Gc preference increased (**Fig. 6-9**). However, SLBR interactions with synthetic sialyl disaccharides did not fully predict binding to authentic plasma glycoproteins, which can have larger, more complex underlying glycan structures. This highlights a secondary role for sialic acid identity relative to other determinants, such as glycan linkage and glycoprotein presentation. The discrepancy between plasma glycoprotein binding and synthetic sialyl disaccharide recognition is unlikely to result from alternative glycoprotein engagement. The only other sialic acid derivative present on human cells is Kdn, which accounts for less than 1% of surface glycans and is rarely incorporated into glycoproteins^49^. Additionally, we are not aware of any literature suggesting that SLBRs could engage non-sialoglycoside glycoproteins.

Together, our data support a model in which Neu5Ac/Neu5Gc preference modulates, but does not uniquely determine, SLBR-mediated host glycoprotein recognition by viridans group streptococci. By uncovering distinct structural and allosteric mechanisms of sialoglycan recognition, this work extends our understanding of SLBR specificity and host range, offering insight into bacterial adaptation. Finally, our work identifies that models of host tropism will benefit from a nuanced view of glycoprotein chemistry that encompasses sialic acid identity, glycan context, and presentation.

### Experimental Procedures

#### Protein Expression and Purification for X-ray Crystallography

SLBR_Hsa_ was expressed and purified as previously described^16^. Briefly, the pSV278 vector (Vanderbilt university) appends a His-maltose-binding protein (MBP) tag at the N-terminus of SLBR_Hsa,_ followed by a thrombin cleavage site. His_6_-MBP-SLBR_Hsa_ was expressed in *Escherichia coli* BL21 (DE3) in Terrific Broth medium supplemented with 50 µg/ml kanamycin at 37°C. At an OD_600_ of 1.0, the temperature was lowered to 24°C, and expression was induced with 1 mM isopropyl β-d-thiogalactopyranoside (IPTG) for 6 hrs. Cells were harvested by centrifugation at 5,000 g for 15 min, washed with 0.1 M Tris-HCl, pH 7.5, and stored at −20°C before purification.

Frozen cells were resuspended in buffer containing 20 – 50 mM Tris-HCl, pH 7.5, 150 – 200 mM NaCl, 1 mM EDTA, 1 mM PMSF, 2 µg/ml Leupeptin, 2 µg/ml Pepstatin, then disrupted by sonication. Lysate was clarified by centrifugation at 38,500 g for 35 – 60 min and passed through a 0.45 µm filter. Purification of His_6_-MBP-SLBR_Hsa_ was performed at 4°C with an MBP-Trap column and eluted in 10 mM maltose. Eluted proteins were concentrated in a 10 kDa molecular weight cutoff concentrator and exchanged into buffer containing 20 mM Tris-HCl, pH 7.5, and 200 mM NaCl. The His_6_-MBP affinity tags were cleaved from SLBR_Hsa_ with 1 U of thrombin per mg of His_6_-MBP-SLBR_Hsa_ overnight at 4°C. The cleaved His_6_-MBP tag was separated from pure SLBR_Hsa_ using a Superdex 200 Increase 10/30 GL column equilibrated in 20 mM Tris-HCl, pH 7.5, and 200 mM NaCl. After purification, the protein was > 95% pure, as assessed by SDS-PAGE, and was stored at −80°C.

#### Crystallization and Collection of X-ray Diffraction Data

Crystals of SLBR_Hsa_ (21.6 mg/ml in 20 mM Tris-HCl, pH 7.2) were formed by the sitting drop vapor diffusion method by equilibrating 1 µL protein and 2 µL reservoir solution over 50 µL of reservoir solution (0.1 M Succinate/Phosphate/Glycine pH 10.0 and 25% PEG 3350). Co-crystals of SLBR_Hsa_ with sialoglycan ligands were prepared by soaking fully formed crystals in reservoir solution supplemented with 5 mM of each ligand for 20 hr. Crystals did not require cryoprotection beyond the reservoir solution and were cryocooled by plunging into liquid nitrogen. Data were collected at −180°C on beamline 9-2 at the Stanford Synchrotron Radiation Lightsource. Data were processed in HKL2000^50^.

#### Structure Determination and Refinement

The structure of each sialoglycan-bound SLBR_Hsa_ was determined using isomorphous replacement by removing all solvent molecules from unliganded SLBR_Hsa_ (PDB entry 6EFC^16^) and performing rigid body refinement in PHENIX^51^. The resultant model was improved using alternate rounds of model building in COOT^52^ and refinement in PHENIX^51^. Throughout the process, the R_free_ reflections were selected to be the same as for the unliganded SLBR_Hsa_ (PDB entry 6EFC^16^). Data collection and refinement statistics are listed in Table 1.

#### Sialoglycan reagents

Neu5Acα2-3Gal and Neu5Gcα2-3GalβOMe used in crystallography studies were prepared as previously reported^32,33^. Biotinylated sialyl disaccharides for the ELISAs were purchased from Sigma (GNZ-0035-BM for Neu5Ac and GNZ-0018-BM for Neu5Gc).

#### MD Simulations

The crystal structures of SLBR_Hsa_ bound to either Neu5Acα2-3Gal or Neu5Gcα2-3Gal were used to generate starting models for MD simulations. MD was performed on the proteins and glycan ligands using the Amber14 ff14SB^53^ and Glycam06^54^ force fields, respectively, with a non-bonded cutoff of 10 Å using the Particle Mesh Ewald algorithm^55^. Each protein-glycan system was hydrated by water model TIP3P^56^ using an octahedral box of 10 Å around the protein in each direction. Initially, the protein was held fixed with a force constant of 500 kcal mol^-1^ Å^-2^ while the system was energy minimized with 500 steps of steepest descent. This was followed by 500 steps of energy minimization with the conjugate gradient method. In a second minimization step, the restraints on the protein were removed, and 1000 steps of steepest descent minimization were performed, followed by 1500 steps of conjugate gradient. The system was heated to 300 K while holding the protein fixed with a force constant of 10 kcal mol-1 Å-2 for 1000 steps. Then, the restraints were removed, and 1000 MD steps were performed. The SHAKE algorithm^57^ was used to constrain all bonds involving hydrogen in the simulations. MD production runs were performed at 300 K using the NPT ensemble and a 2-fs time step. The temperature was fixed with the Langevin dynamics thermostat^58^, and the pressure was fixed with the Monte Carlo barostat^59^. Three independent runs were performed for each simulation. All analyses were done using the Pytraj package^60^.

#### Protein Expression and Purification for ELISAs and Far Western Blotting

SLBR_Hsa_, SLBR_SrpA_, and all variants were expressed and purified as previously described^16^. Point mutants were created from already cloned SLBRs in vector pBG101 (Vanderbilt University), which encodes a His_6_-Glutathione-S-transferase (GST) tag at the N-terminus, followed by a 3C protease cleavage site. His_6_-GST-SLBRs were expressed in Escherichia coli BL21 (DE3) in Miller’s Luria Broth at 37°C with 50ug/mL of kanamycin for ∼3 hr to reach an A_600_ of 0.80. SLBR expression was induced with 1 mM IPTG for 3 hours at 24°C. Cells were harvested by centrifugation at 7,000 x g for 20 minutes. Cell pellets were resuspended in 125 mM Tris, 150 mM sodium chloride, pH 8.0, supplemented with 1 mM EDTA, 1 mM PMSF, 2 µg/ml Leupeptin, 2 µg/ml Pepstatin, then lysed by sonication. Lysate was clarified by centrifugation at 18,000 x g for 1 hour. The supernatant was filtered (0.45µm) and purified using glutathione-sepharose as instructed by the manufacturer (Thermo, 16108).

After purification, proteins were concentrated in a 30 kDa molecular weight cutoff concentrator and buffer exchanged using a Superdex 200 Increase 10/30 GL column equilibrated in 1X Dulbecco′s Phosphate Buffered Saline (DPBS) with calcium and magnesium (Sigma, D1283). The protein was > 95% pure, as assessed by SDS-PAGE, and was stored at −80°C.

#### ELISAs

The binding of biotinylated sialoglycans to immobilized GST-SLBRs was performed as described^2,7,16^. In short, purified SLBRs were diluted to 500 µM in DPBS and added to a 96-well microtiter plate. Plates were incubated overnight at 4°C. Unbound proteins were removed by aspiration, and wells were rinsed with DPBS. Biotinylated glycans were diluted to the indicated concentrations in DPBS containing 1X Blocking Reagent (Roche, 11585762001) and incubated for 1 hr at room temperature. Wells were rinsed three times with DPBS. Streptavidin-conjugated horseradish peroxidase (Sigma, S5512) was added to each well, and the plate was incubated for 1 hr at room temperature. The wells were washed twice with DPBS, and then a solution of 0.4 mg o-Phenylenediamine dihydrochloride (Sigma, P8787) per mL phosphate-citrate buffer (Sigma, P4922) was added to the wells. The absorbance at 450 nm was measured after approximately 20 min. Data were plotted as the means ± standard deviations, with n = 3.

#### Far-Western Blotting

Far western blotting was previously described^2,7,16^. Briefly, human or rat plasma (Innovative Research) was diluted 1:10 into 10 mM Tris, 1 mM EDTA, pH 8, combined with LDS sample buffer and DTT (50 mM final concentration). Samples were boiled for 10 min, and proteins were separated by electrophoresis on 3–8% polyacrylamide gradient gels (Life Technologies), and then transferred to BioTraceNT (Pall Corporation). Membranes were incubated for 1 h at room temperature with 1× Blocking Reagent in DPBS. GST-SLBRs were then added to a final concentration of 5 nM, and the membranes were incubated for 90 min at room temperature with gentle rocking. After rinsing three times with DPBS, the membranes were incubated for 1 h at room temperature with anti-GST diluted 1:5000 in DPBS containing 1× Blocking Reagent. Membranes were rinsed three times with DPBS and then incubated for 1 h at room temperature with horseradish peroxidase-conjugated goat anti-rabbit antibodies diluted 1:50,000 in DPBS. Membranes were again rinsed three times with DPBS and then developed with SuperSignal West Pico (Thermo Scientific).

## Acknowledgements

We thank L. Loukachevitch for experimental assistance in the early stages of this work. This work was supported by NIH grant GM137458 to TMI, BAB, XC, JCS, and DE019807 to SR, TMI. KMM was supported by NIH training grant GM007628. HES was supported by the NIH Training grant EY007135. Use of the Stanford Synchrotron Radiation Lightsource, SLAC National Accelerator Laboratory, is supported by the U.S. Department of Energy, Office of Science, Office of Basic Energy Sciences under Contract No. DE-AC02-76SF00515. The SSRL Structural Molecular Biology Program is supported by the DOE Office of Biological and Environmental Research, and by the National Institutes of Health, National Institute of General Medical Sciences (P30GM133894). The contents of this publication are solely the responsibility of the authors and do not necessarily represent the official views of NIGMS or NIH.

