## Supplementary Information for "Sialic Acid Identity Modulates Host Tropism of Sialoglycan-binding Viridans Group Streptococci"

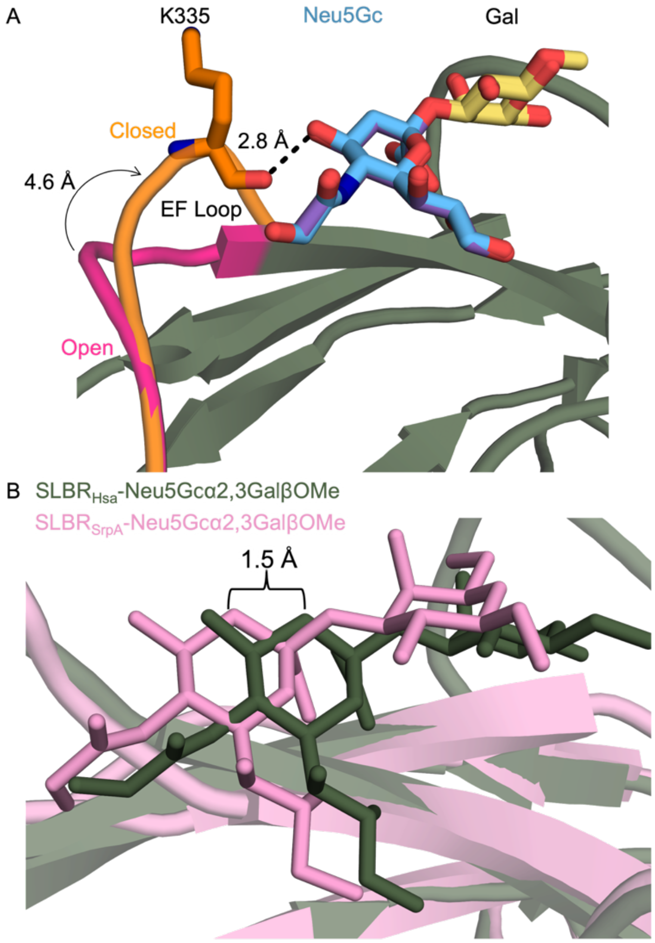


**Supplementary Figure 1 Aspects of ligand binding in $SLBR_{Hsa}$ and $SLBR_{SrpA}$. A)** In $SLBR_{Hsa}$-Neu5Gc, the EF loop (*orange*) adopts a closed conformation that allows for a hydrogen bond between the $SLBR_{Hsa}^{K335}$ carbonyl and O4 of Neu5Gc. In $SLBR_{Hsa}$-Neu5Ac, the EF loop (*hot pink*) remains in the open position. **B)** $SLBR_{SrpA}$[1] and $SLBR_{Hsa}$ each bind glycan above the F strand, however there is a lateral shift in position of 1.5 Å with respect to the ΦTRX motif. Shown is each SLBR–Neu5Gc costructure overlaid based on Cα atom positions.

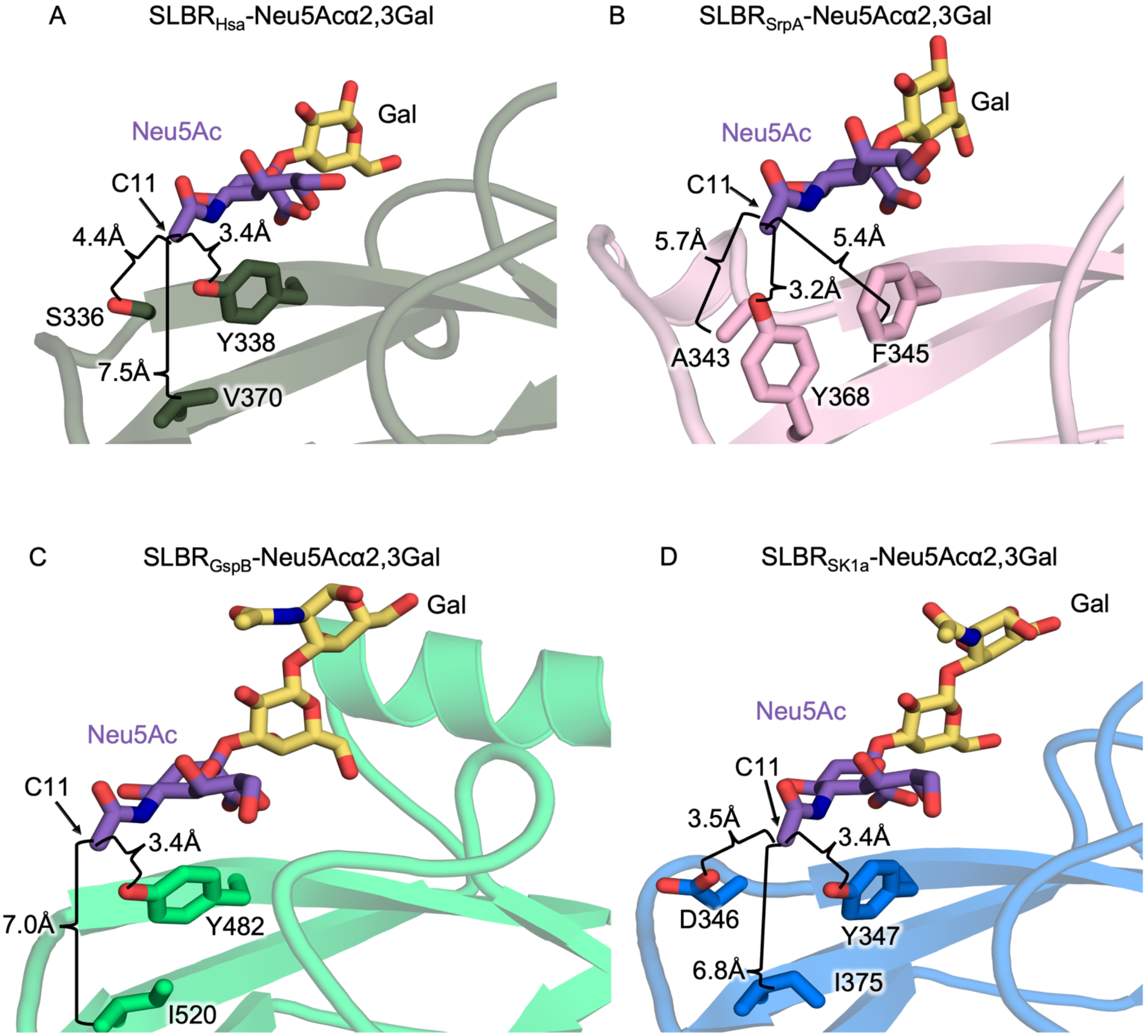


**Supplementary Figure 2 Comparison of Neu5Ac-bound SLBRs. A)** $SLBR_{Hsa}$, **B)** $SLBR_{SrpA}$[1], **C)** $SLBR_{GspB}$[2], and **D)** $SLBR_{SK1a}$[3]. Distances between adjacent non-bonding atoms are shown with brackets.

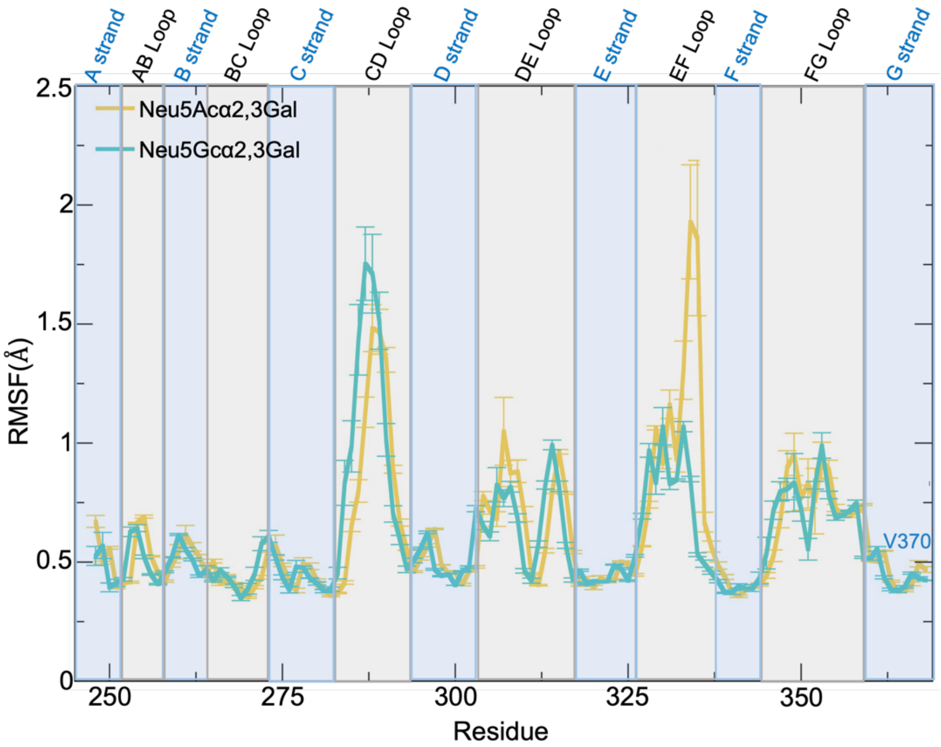


**Supplementary Figure 3 RMSF of the SLBR$_{Hsa}$ bound to Neu5Acα2-3Gal or Neu5Gcα2-3Gal.** RMSF plot for SLBR$_{Hsa}$ residues during MD simulations show backbone flexibility in the presence of Neu5Ac (gold) or Neu5Gc (*cyan*). Sequence regions corresponding to the CD-, EF-, and FG-loops are highlighted with *grey boxes*, while the secondary structural elements are highlighted with *blue boxes*.

A

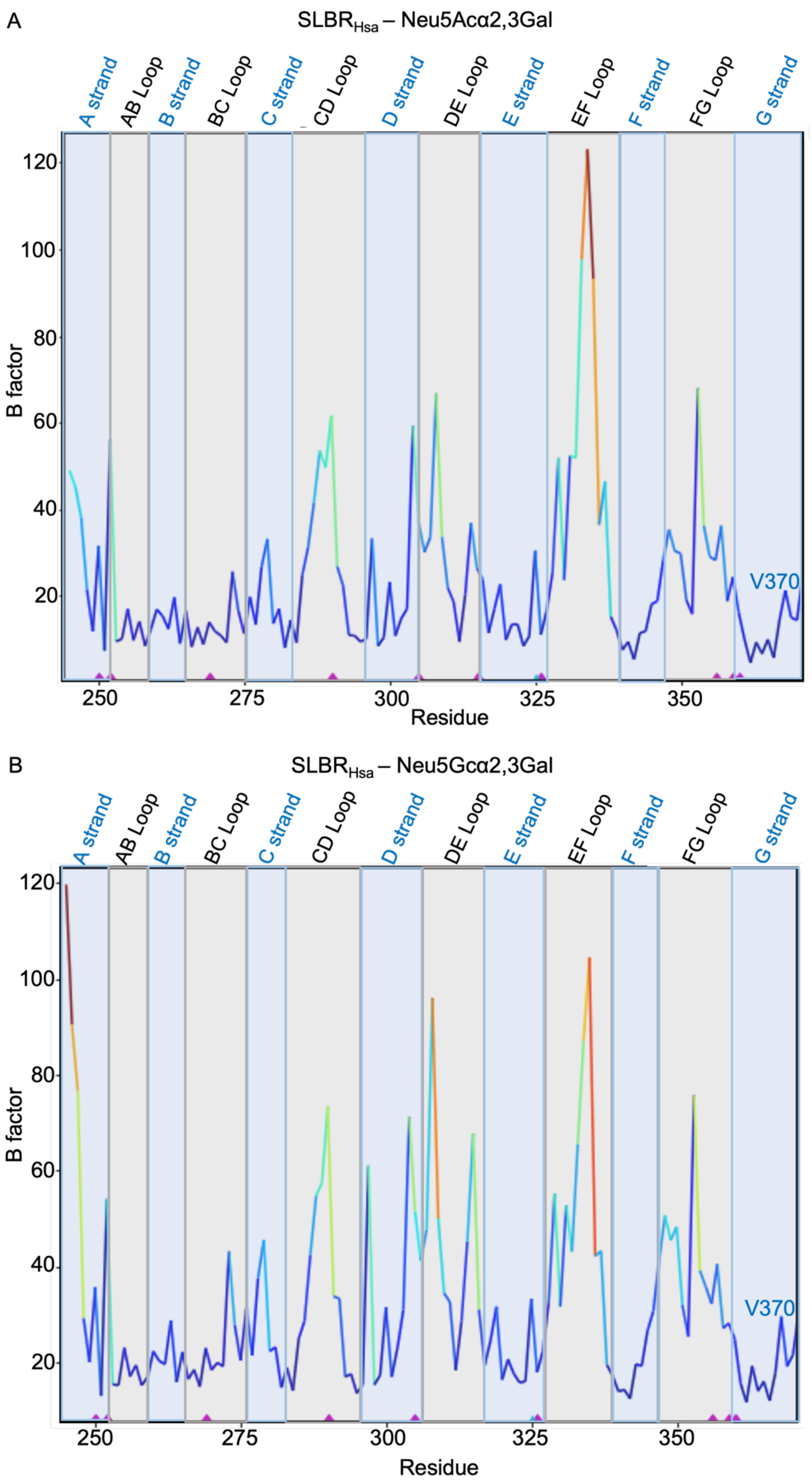


B



**Supplementary Figure 4 Crystallographic temperature factors of SLBR$_{\mathbf{Hsa}}$. A)** Neu5Ac and **B)** Neu5Gc. The temperature factors of the CD-, EF-, and FG-loops are highlighted with *grey boxes*. The temperature factors of the secondary structural elements are highlighted with *blue boxes*. Residues are colored by B-factor magnitude, with warmer colors representing a higher temperature factor.

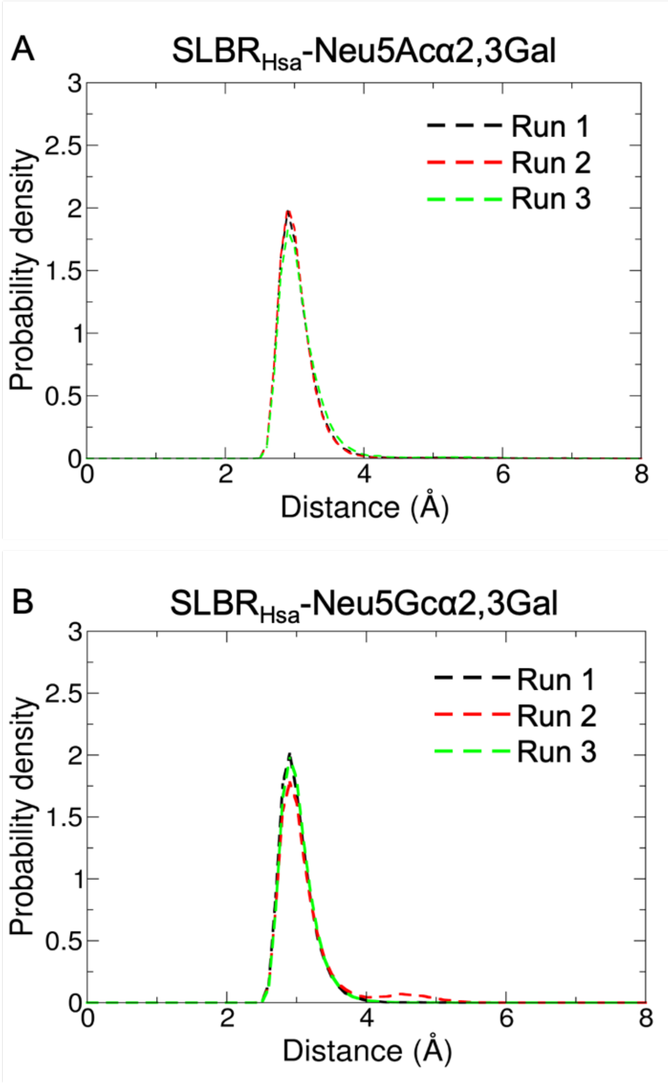


**Supplementary Figure 5 Probability distributions of the distance of the EF-loop in SLBR$_{Hsa}$.** Distance between SLBR$_{Hsa}^{K335}$ backbone carbonyl and Neu5Gc-O4 in simulations of SLBR$_{Hsa}$ bound to disaccharides terminated in **A)** Neu5Ac and **B)** Neu5Gc. The ~3 Å distance is consistent with a hydrogen-bonding interaction that accompanies EF loop closure of the ligand for the duration of the simulation. All simulations were performed in triplicate.

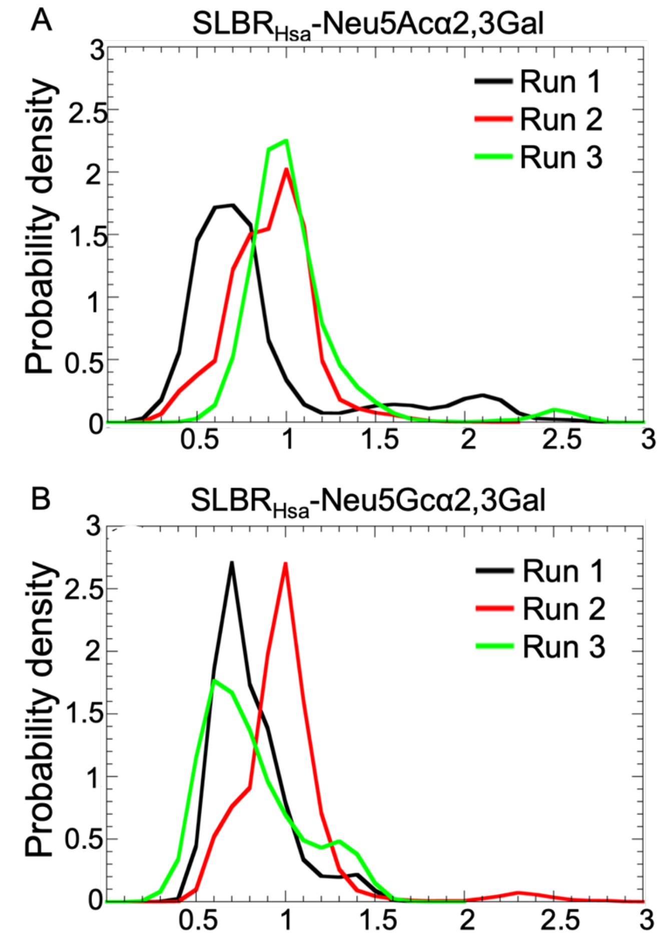


**Supplementary Figure 6 Probability distribution RMSD of disaccharides bound to SLBR$_{Hsa}$. A)** Neu5Ac and **B)** Neu5Gc. Three independent molecular dynamics simulations were performed. Mean RMSD Neu5Ac = 0.85 Å (±0.14 Å). Mean RMSD Neu5Gc = 0.83 Å (±0.15 Å).

### Supplementary Information References

1 Bensing, B. A. *et al.* Structural Basis for Sialoglycan Binding by the Streptococcus sanguinis SrpA Adhesin. *J Biol Chem* **291**, 7230-7240, doi:10.1074/jbc.M115.701425 (2016).

2 Pyburn, T. M. *et al.* A structural model for binding of the serine-rich repeat adhesin GspB to host carbohydrate receptors. *PLoS Pathog* **7**, e1002112, doi:10.1371/journal.ppat.1002112 (2011).

3 Stubbs, H. E. *et al.* Tandem sialoglycan-binding modules in a Streptococcus sanguinis serine-rich repeat adhesin create target dependent avidity effects. *J Biol Chem* **295**, 14737-14749, doi:10.1074/jbc.RA120.014177 (2020).
